# Single-Cell Proteomics Defines the Cellular Heterogeneity of Localized Prostate Cancer

**DOI:** 10.1101/2021.01.25.428046

**Authors:** Laura De Vargas Roditi, Andrea Jacobs, Jan H. Rueschoff, Pete Bankhead, Stephane Chevrier, Hartland W. Jackson, Thomas Hermanns, Christian D. Fankhauser, Cedric Poyet, Felix Chun, Niels J. Rupp, Alexandra Tschaebunin, Bernd Bodenmiller, Peter J. Wild

## Abstract

Localized prostate cancer exhibits multiple genomic alterations and heterogeneity at the proteomic level. Single-cell technologies capture important cell-to-cell variability responsible for heterogeneity in biomarker expression that may be overlooked when molecular alterations are based on bulk tissue samples. The aim of this study was to identify novel prognostic biomarkers and describe the heterogeneity of prostate cancer and the associated immune cell infiltrates by simultaneously quantifying 36 proteins using single-cell mass cytometry analysis of over 1,6 million cells from 58 men with localized prostate cancer. To perform this task, we proposed a novel computational pipeline, Franken, which showed unprecedented combination of performance, sensitivity and scalability for high dimensional clustering compared to state of the art methods. We were able to describe subpopulations of immune, stromal, and prostate cells, including unique changes occurring in tumor tissues and high grade disease providing insights into the coordinated progression of prostate cancer. Our results further indicated that men with localized disease already harbor rare subpopulations that typically occur in castration-resistant and metastatic disease, which were confirmed through imaging. Our methodology could be used to discover novel prognostic biomarkers to personalize treatment and improve outcomes.

## INTRODUCTION

The treatment of localized prostate cancer is based on clinicopathological information including Gleason score, prostate-specific antigen (PSA) levels, stage and patient age^1^. While the majority of patients with localized disease can be cured, some men recur with metastatic disease^2^ because of microscopic spread. The observed heterogeneity of outcomes might be explained by heterogeneity within tumors^3^, which is missed by the current grading system and new prognostic biomarkers are of utmost importance.

Several potential biomarkers including gene fusions, mutations, epigenetic heterogeneity, and proteins have been studied^4^. Technological advances in proteomics now allow both exploration of the proteome for biomarkers and assessment of the heterogeneity of biomarker expression. However, analysis of a whole tissue core misses important cell-to-cell variability. In ths study, we performed mass cytometry analysis of dissociated single cells from prostatectomies of 58 patients with tumors at varying grades and UICC (Union internationale contre le cancer) stages using a set of 36 metal-tagged antibodies that recognize surface markers, enzymes, transcription factors, and markers of functional readouts selected to facilitate characterization of the phenotypic diversity of prostate tumors and their microenvironment. The power to comprehensively analyze heterogeneity of tumors by simultaneously measuring dozens of markers in hundreds of thousands to millions of cells makes mass cytometry the ideal tool to characterize single-cell subpopulations present in prostate or other tumors including those rare populations that cannot be detected with lower parametricity or lower throughput methods.

Although mass cytometry has single-cell resolution capabilities, there are statistical challenges involved in analyzing such high-dimensional data. State-of-the-art clustering methods either underperform in precision and recall or require unnecessarily long runtimes and prohibitive computational resources^5^. To address this issue we developed an unsupervised, single-cell clustering approach, Franken, which is unmatched in its combination of speed and performance. Use of Franken to quantify the phenotypic diversity of single cells in prostate tumor samples identified unique progression-related single-cell phenotypes. We detected immune landscape features unique to patients with high grade prostate cancer, reflected by higher frequencies of macrophage and T cell phenotypes than observed in patients with intermediate grade disease. Further, we observed tumor-specific prostate epithelial phenotypes, including AR-negative and/or PSA-negative phenotypes typically associated with resistance to ADT and CRPC^6-8^, and previously undescribed and rare CD15^+^ phenotypes.

## RESULTS

### Clustering of high-dimensional mass cytometry data defines molecular profiles of prostate subpopulations

Using mass cytometry, we profiled tumor samples from 58 prostate cancer patients, including 24 patients with the International Society of Urological Pathology (ISUP) grade II (Gleason score 3+4), 22 grade III cases (Gleason score 4+3), and 12 patients with grade V prostate carcinomas (Gleason scores 4+5, 5+4 or 5+5) (Figure 1a). Fresh tissue samples for dissociation into single cell suspensions were collected from 58 prostate cancer patients. From 41 of 58 (71%) patients, the tumor could not be demarcated macroscopically. The presence of prostate cancer was confirmed histologically after examining the opposite side of the specimen, first in the frozen section and then after paraffin embedding; these samples are referred to as random prostate tissue (RPT) samples. For a further 17 patients (29%), paired samples were taken from a macroscopically visible tumor mass and adjacent benign prostatic tissue (ABPT); tumor samples were validated using a frozen section of the opposite side. ABPT specimens were taken from the contralateral transitional zone of the prostate and never from the peripheral one, where tumor is more likely to be located. Single-cell suspensions from all prostate tissue samples and from 10 cell lines, including prostate cancer, stromal, and immune cells (Supplemental table 1), were barcoded, pooled and stained with a 36-antibody prostate cancer-centric panel, before mass cytometry acquisition. The antibody panel was designed to quantify markers that identify prostate epithelial cells, cells of the stroma and immune microenvironment, and markers of proliferation and survival (Figure 1b). Data for a total of 1,670,117 live cells were generated.

**Figure 1.**
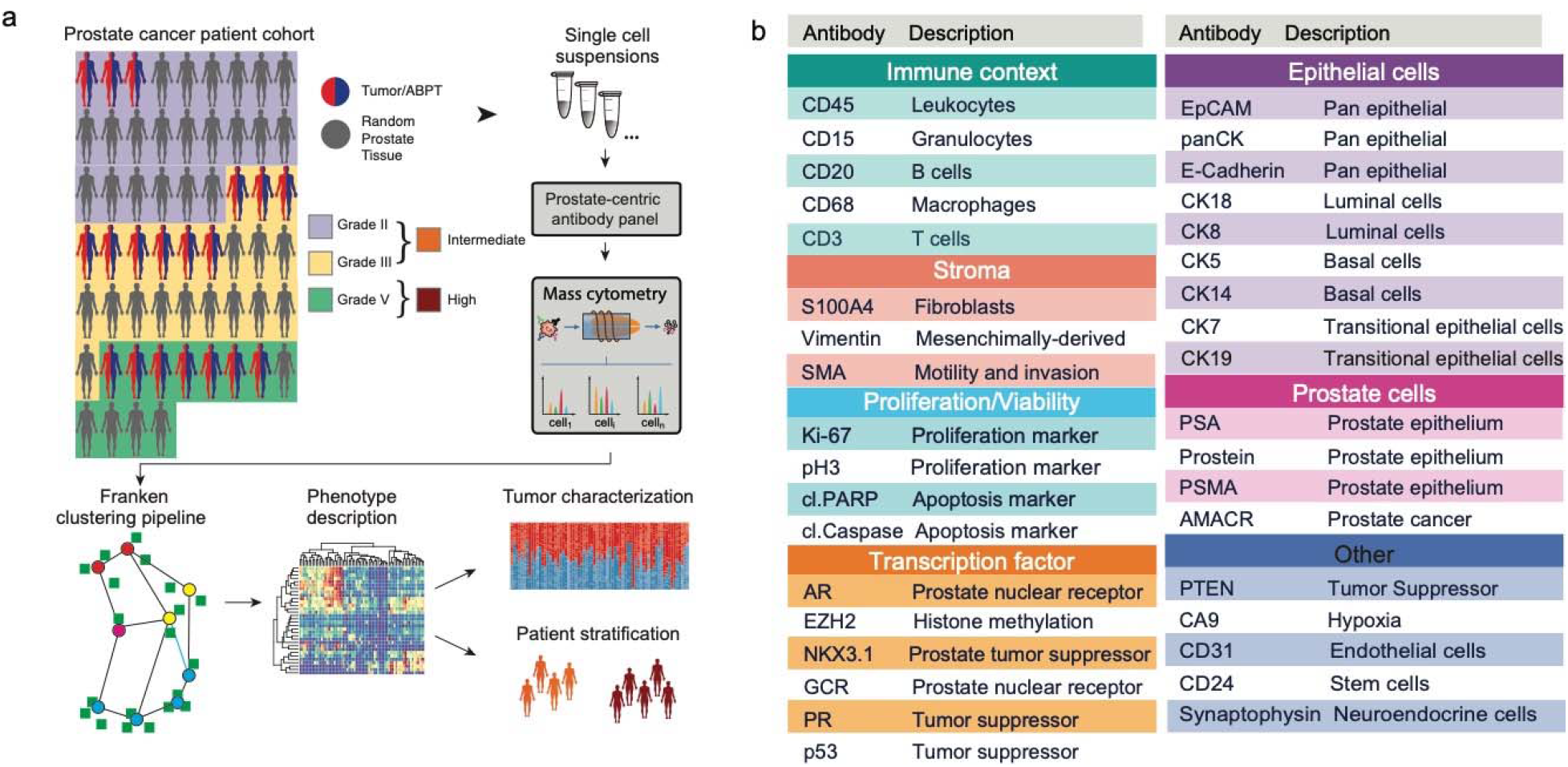
Schematic of method for characterization of primary prostate cancer tissue using mass cytometry. (a) The patient cohort consisted of 58 primary prostate cancer cases. For 16 patients, tumor and adjacent benign prostatic tissue (ABPT) samples were available. The remaining samples were from randomly selected regions from a prostatectomy without tumor assessment. Samples were analyzed by mass cytometry, and data were analyzed using Franken. (b) Markers used to categorize prostate epithelial cells as luminal, basal, or transitional, and markers used to identify tumor cells, cells from the microenvironment and functional features, such as proliferation, apoptosis or hypoxia.

The high dimensionality of this dataset represented a challenge for data visualization and clustering. To address this task, we developed a highly efficient computational pipeline called Franken (Supplemental Figure 1a). The pipeline begins by building a large self-organizing map (SOM), which is used to fit all of the data. The SOM nodes in the original high-dimensional space are then used to build a mutual nearest neighbor graph using the Tanimoto similarity^9^ (also known as extended Jaccard similarity) that is subsequently clustered via the Walktrap graph clustering technique^10^. Although all results presented in this paper were obtained in an unsupervised manner, the pipeline also offers the option to define the chosen number of clusters (detailed description found in methods). We demonstrated the F1 performance and scalability of Franken when compared to state-of-the-art methods on two independent mid-size CyTOF datasets (around 200,000 cells each; Supplemental Figure 1b-d). We also showed that our pipeline is robust to its parameters’ choice and scalable up to dozens of millions of cells in a synthetic dataset (Supplemental Figure 1e,f).

Franken’s scalability and ability to resolve rare metaclusters made it uniquely suitable to explore our new prostate cancer patient dataset containing 1,670,117 cells. Analysis of the prostate cancer dataset using Franken identified a total of 55 clusters (Figure 2a). Franken is a very sensitive technique and leads to fine grained clusters that may represent the gradient expression of certain markers without clear biological differences. It must be noted that discrete labels do not easily apply to datasets with continuous expression such as single-cell data, where a clear cutoff between cell states does not necessarily exist. Nonetheless, to obtain clusters that were qualitatively different in terms of marker combination (which markers were expressed, instead of how much), Franken clusters were further merged into 33 metaclusters using hierarchical clustering of Pearson correlation dissimilarities with average linkage. We found 14 epithelial, 16 immune, one stromal and one endothelial phenotypes, based on marker expression profiles (Supplemental figure 2a). We also identified one cluster which was mostly negative for all 36 markers in the panel (denoted as NE01). Given that our panel does not cover the entire proteome, this may represent a cell type not characterized by the markers in our panel or simply outliers and was excluded from further analysis. All metaclusters were annotated using a two-letter and two-digit identifier ranked by decreasing metacluster size (TC01 > TC02 > TC03 >&) for each cell category. The total number of cells in a cluster ranged from a few hundred (437 cells in EP01) to hundreds of thousands (391,554 in TC01; Supplemental Figure 2a).

**Figure 2.**
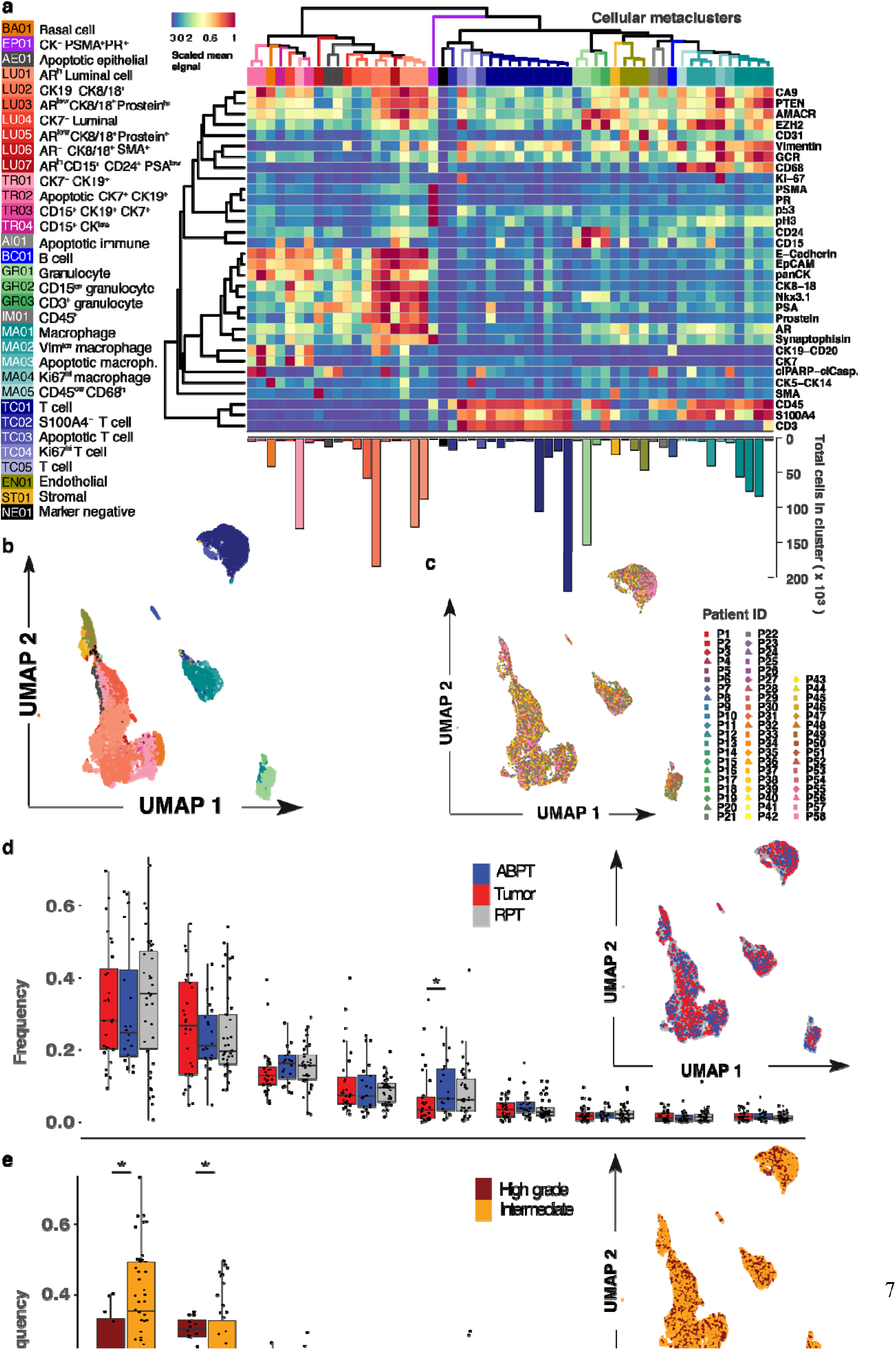
Prostate cancer samples have similar overlapping phenotypic profiles. (a) Heatmap of scaled mean signal of marker expression in 55 Franken clusters; numbers are colored according to metaclusters resulting from hierarchical clustering merging (using Pearson correlation dissimilarities) of Franken clusters. Bar plot below the heatmap corresponds to the number of cells found in each cluster. (b) UMAP map of 23,200 (400 per patient) cells colored by cellular metacluster as indicated in panel (a). (c) UMAP map of 23,200 (400 per patient) cells colored by patient. (d) (e) Boxplots of frequencies of the main cell types across all 58 samples from tumor, ABPT, and RPT. Significant changes were seen between tumor and ABPT in the proportion of granulocytes (two-sided Wilcoxon signed rank paired test, p = 0.008). N=17. (e) Boxplots of frequencies of the main cell types in samples from all 58 patients in our cohort stratified by intermediate and high grade tumors. Changes in luminal and T cell compartments are significant according to a two-sided Mann-Whitney-Wilcoxon test (p = 0.028 and 0.014, respectively). Intermediate N = 46 and high grade N = 12.

In order to project the high dimensional data into a two-dimensional representation, we used the UMAP (Uniform Manifold Approximation and Projection) method for dimensionality reduction visualization^29^. UMAP showed that our analysis recapitulated the main cell-type compartments within the prostate (Figure 2b); also detected were rarer cell states such as apoptotic cells (TR02, AE01, AI01, MA03 and TC03). Franken was capable of resolving very rare populations present at frequencies as low as 1/5,000 (PR-high metacluster EP01; Supplemental Figure 2b). FlowSOM, although very efficient in terms of speed, revealed fewer and larger subpopulations than Franken; very rare populations identified by Franken could not be identified by FlowSOM or PhenoGraph. Franken, therefore, provides an unprecedented combination of performance, sensitivity and speed compared to existing clustering methods^19^. To ensure good separability of each class (metacluster) and quality of the final clustering configuration, we trained a logistic regression classifier with lasso regularization (using 5-fold cross-validation to identify the regularization parameter λ; Supplemental Figure 2c,d). Most metaclusters could be predicted with higher than 99% accuracy, and the lowest at 93%.

All detected clusters contained cells from ABPT and tumor regions (Supplemental Figure 3a). This suggested that tumor cells were present within the ABPT tissue and/or that ABPT tissue was present inside the tumor mass which was expected due to the usual way prostate tumors spread in the prostate as well as intrinsic limitations of the macroscopic-based sample collection procedure (which could not ensure the adjacent regions were 100% tumor-free). Alternatively, this could suggest our custom panel missed important markers to make such distinction.

Luminal cells were the most abundant cell type in the prostate and corresponded on average (across all 58 patients) to 32% of a patient’s sample; T cells were the second most abundant population (24% on average). When combined, cells from the immune compartment and other cells of the tumor microenvironment made up over half the cells (54% on average) found within samples of this cohort (Supplemental Figure 3b). Franken clustering identified a range of prostate epithelial phenotypes including a single basal cell phenotype characterized by CK5 and CK14 expression (BA01), four transitional epithelial phenotypes expressing a combination of CK7 and CK19 (TR01-04), and seven epithelial luminal phenotypes (LU01-07) defined by the expressed CK8, CK18. Cellular metaclusters that contained a combination of CK7 or CK19 and CK8 or CK18 were annotated as luminal epithelial cells. Only CK7 and CK19-positive metaclusters with very low to no CK8 and CK18 expression were denoted as transitional epithelial cells. Luminal epithelial cells also expressed a combination (or varying expression intensities) of AR, PSA, Prostein, Synaptophysin, AMACR, EZH2, PTEN and Nkx3.1. These were weakly or not expressed in transitional or basal cell metaclusters.

In the microenvironment, five different T cell phenotypes were detected expressing CD3 and CD45, and five macrophage phenotypes were characterized by CD68 and CD45 expression. Also detected were three granulocyte (expressing CD24 and/or CD15), one stromal (characterized by SMA and S100A4), one B cell (CD20-expressing), and one endothelial (CD31-expressing) clusters. Unlike in previous works^11^, we did not observe patient-specific batch effects, which could have led to each patient clustering separately from one another. We found that each metacluster contained a mixture of cells from most patients as illustrated in the UMAP visualization (Figure 2c).

There was considerable overlap in the single-cell phenotypes present within paired tumor and ABPT regions (UMAP; Figure 2d). Samples were stratified according to tumor, ABPT and RPT across cell types and we found significantly lower frequency of granulocytes in tumor regions than in ABPT (Figure 4e. Visualization of cells from intermediate grade (ISUP grades II and III) and high grade tumors (ISUP grade V) also revealed significant overlap (UMAP; Figure 2e) and indicated lower frequencies of luminal cells and higher frequencies of T cells in samples from patients with high grade compared to intermediate grade disease (Figure 2d).

### Immune landscape differs between tumor and benign adjacent tissue and across prostate cancer ISUP grade

The UMAP visualization of 23,200 (400 per patient) cells, randomly selected across the patient cohort, revealed the expression patterns of markers associated with the microenvironment (Figure 3a and Supplemental Figure 3c). We validated both observations by quantifying T cells (CD3^+^) and granulocytes (CD15^+^) in a tissue microarray (Supplemental Figure 3d and data availability section) that included formalin-fixed paraffin-embedded tissues from all patients in the cohort. Confirming the mass cytometry data, we observed higher densities of T cells in high grade tumors (Figure 3b and c) than in intermediate grade prostate tumors and lower densities of granulocytes in tumor regions than in ABPT regions (Figure 3d and 3e).

**Figure 3.**
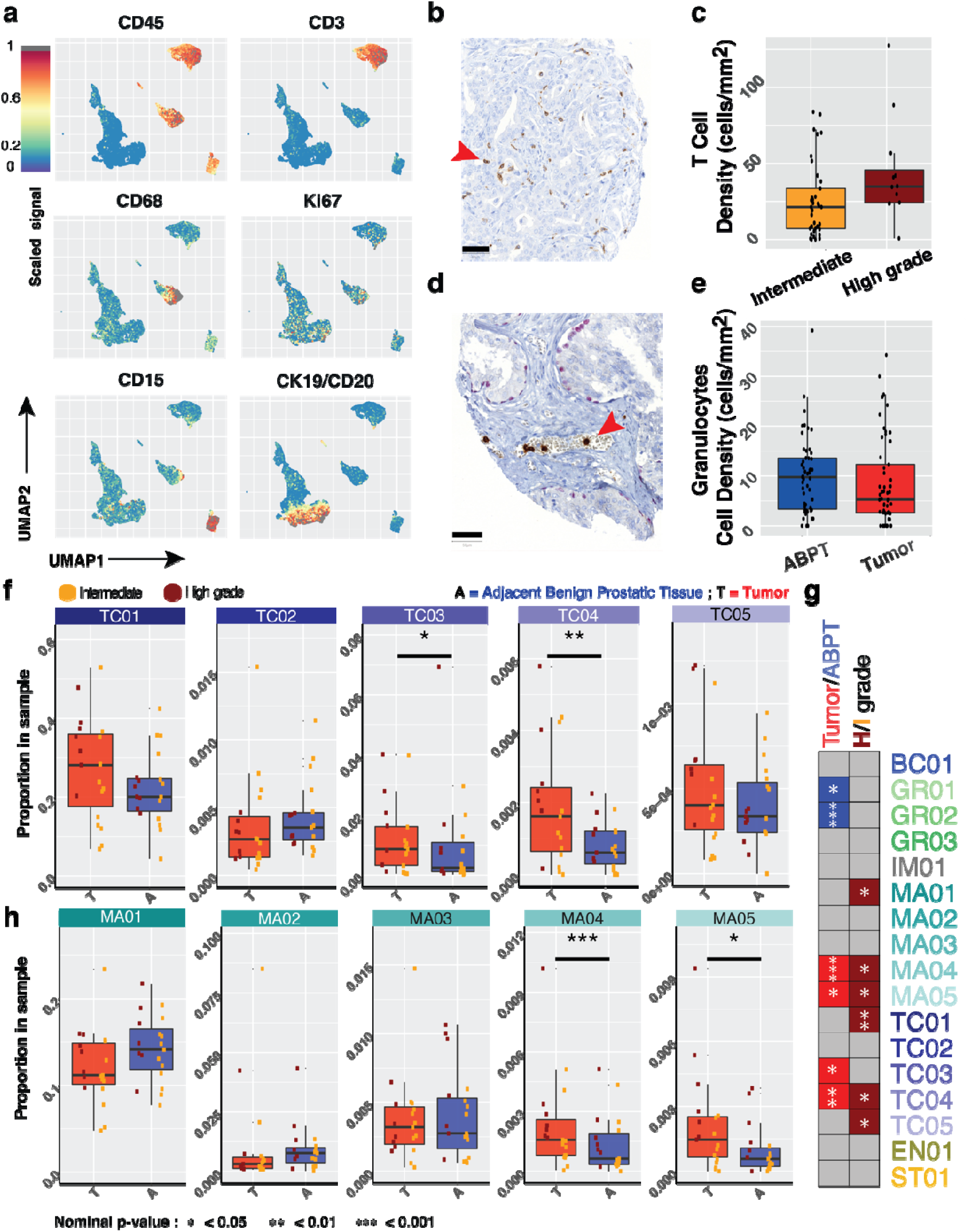
Stratification of samples reveals prostate tissue changes associated with tumor and advanced disease. (a) UMAP of 23200 cells (400 per patient) colored by expression of indicated marker (b) Representative tissue sample stained for CD3 from a tissue microarray generated from prostate samples from the same cohort analyzed by mass cytometry. (c) Densities of T cells as determined by CD3 staining (p = 0.042). (d) Representative tissue sample stained for CD15 from a tissue microarray generated from prostate samples from the same cohort analyzed by mass cytometry. (e) Densities of granulocytes as determined by CD15 staining (p = 0.058). Scale bar, 50 μm. (f) Proportion of T cell metaclusters in ABPT and tumor samples across patients with paired samples (p = 0.066, 0.169, 0.023, 0.002 and 0.332 for TC01-05, respectively). N=17. (g) Summary table of clusters that were significant when comparing ABPT and Tumor samples (N=17 for both groups). Metaclusters enriched in ABPT are colored in blue while those enriched in tumor samples are colored in red. Comparison between intermediate and high grade patient samples (for combined tumor/ABPT; Intermediate N = 46 and high grade N = 12). Metaclusters enriched in patients with high grade disease are colored in dark red. Only significant relationships are colored and remaining comparisons are shown in grey. (h) Proportion of macrophage metaclusters in ABPT and tumor samples across patients with paired samples (p = 0.095, 0.169, 0.515, 0.0004 and 0.014 for MA01-05, respectively). N=17. In panels (f) and (h) dots are colored by disease severity (intermediate vs high grade). In all boxplots, boxes illustrate the interquartile range (25th to 75th percentile), the median is shown as the middle band, and the whiskers extend to 1.5 times the interquartile range from the top (or bottom) of the box to the furthest datum within that distance. Statistical testing between dependent paired Tumor and ABPT samples was done using a Wilcoxon signed rank paired-sample statistical tests (two-sided). Independent intermediate and high grade samples were tested using a two-sided Wilcoxon rank sum test. Number of patients in each group is indicated by N.

We identified multiple subpopulations of each immune cell type. By focusing on the comparison of each individual metacluster, we found that two T cell clusters were significantly enriched in tumor samples (TC03 and TC04, apoptotic T cells and proliferating T cells, respectively) when compared to the adjacent ABPT regions (Figure 3f). Notably, proliferating T cells were also enriched in high grade tumors and in high grade patient samples when tumor and adjacent tissue were mixed (Figure 3g and Supplemental Figure 4a) suggesting that this T cell phenotype is enriched throughout the prostate of patients with high grade disease and not only in the core of the tumor.

The overall frequencies of macrophages were not significantly different between tumor and tumor-adjacent (ABPT) samples (Figure 2d). However, two macrophage metaclusters were enriched in tumor samples (MA04 and MA05, proliferating macrophages and CD45^low^ macrophages respectively). These same metaclusters were further enriched in high grade patient samples (Supplemental Figure 4b). The overall macrophage proportion was actually lower in tumor samples than in tumor-adjacent samples, highlighting the importance of analyzing such a complex dataset at single-cell resolution to reveal that rare macrophages phenotypes can change in the opposite trend to the overall macrophage population. The majority of macrophages are localized in the prostate stroma, but their density is greater in tumorigenic regions^12,13^. This is a confounding factor when comparing macrophage frequencies across tumor grades, since lower grade tumors have a greater proportion of stroma than high grade tumors, resulting in a higher frequency of stroma-infiltrating macrophages (Figure 3h).

In summary, our clustering analysis identified changes in the cellular phenotypes present in the prostate tumor microenvironment compared to adjacent ABPT regions. Distinct macrophage phenotypes were associated with prostate tumors and with the stroma rich ABPT regions. Overall, the cell-type compositions of the tumor microenvironments differed with tumor grade, with the exception of granulocytes, which were decreased in tumor regions regardless of grade (Supplemental Figure 4c). Although the tumor microenvironment of the intermediate sub-cohort was characterized mostly by a relative decrease of immune phenotypes compared with ABPT regions, we found the opposite in the high grade sub-cohort where multiple immune phenotypes were enriched, notably highly proliferative macrophage and T cell phenotypes.

### Malignant and benign prostate tissues diverge in rare phenotypes

Matched tumor and ABPT samples exhibited overall similar single-cell phenotypic profiles, and phenotypic profiles were similar across patients (Supplemental Figure 5b). These similarities were likely due to the presence of benign tissue in both tumor and ABPT samples, whereas patient-specific phenotypes are related to heterogeneous, deregulated malignant cells^14^. We found that every sample from every patient, including tumor and ABPT samples, contained basal cells (BA01) as well as CK7^+^/CK19^+^ live and apoptotic transitional epithelial cells (TR01 and TR02, respectively). All patient samples also contained a variety of luminal epithelial cells (LU01-LU07) containing varying combinations of luminal markers CK8/18, AR, PSA, Prostein, Nkx3.1 and in some cases the co-expression of CK19 and CK7 (LU02 and LU04, respectively). However, cell types co-expressing both, CK7 and CK19, expressed little to no CK8/18 (TR01-04). Stem cell marker CD24 and neuroendocrine marker Synaptophysin showed highest expression in luminal epithelial cells.

We carried out statistical comparisons for epithelial metaclusters and summarized the significant relative enrichment results across all metaclusters (Figure 4a and Table S1) for comparisons between patient grade groups (top row; N_I_ = 46, N_H_ = 12), tumor and ABPT (middle row; N_I,H_ = 17), and high versus intermediate grade tumor regions only (bottom row; N_I_ = 10, N_H_ = 7). We identified the enrichment of apoptotic epithelial cells in tumor versus ABPT regions, which was irrespective of tumor grade (Figure 4a and Supplemental Figure 5a). We also found that luminal metaclusters were typically enriched in ABPT regions and/or in intermediate stage patient samples. In particular, Prostein-high and AR-low metaclusters (LU03 and LU05; p values = 0.0001 and 0.026 respectively) were depleted in tumors versus ABPT samples (Figure 4a and Supplemental Figure 5c,d). The depletion of Prostein-high phenotypes was even more pronounced in high grade compared to intermediate tumors (Figure 4a; bottom row). It is possible that during tumor progression, regulation of differentiation programs is lost, and prostate-specific antigens are no longer expressed, supporting the hypothesis that aggressive tumor cells are de-differentiated. A rare SMA-positive luminal cell type (LU06; p value = 0.012) was characteristic of patients with intermediate disease, irrespective of tumor or ABPT region. The only luminal metacluster enriched in high grade patients was a rare PSA low, CD15+, CD24+ (an adhesion protein previously identified as a cancer stem-cell marker^15^) and AR-high cell type (LU07; p value = 0.028; Figure 4a and Supplemental Figure 6). Two additional CD15+ cell populations were identified, TR03 and TR04, amongst transitional epithelial metaclusters. Both were increased in tumor and high grade patient samples, though significant enrichment could only be detected in TR03 (p value = 0.003) which co-expressed CK19 and CK7 (Figure 4a, Supplemental Figure 5e,f) while TR04 may be a more common precursor with lower cytokeratin expression. TR03 and TR04 also expressed a low amount of basal markers CK5/14.

**Figure 4.**
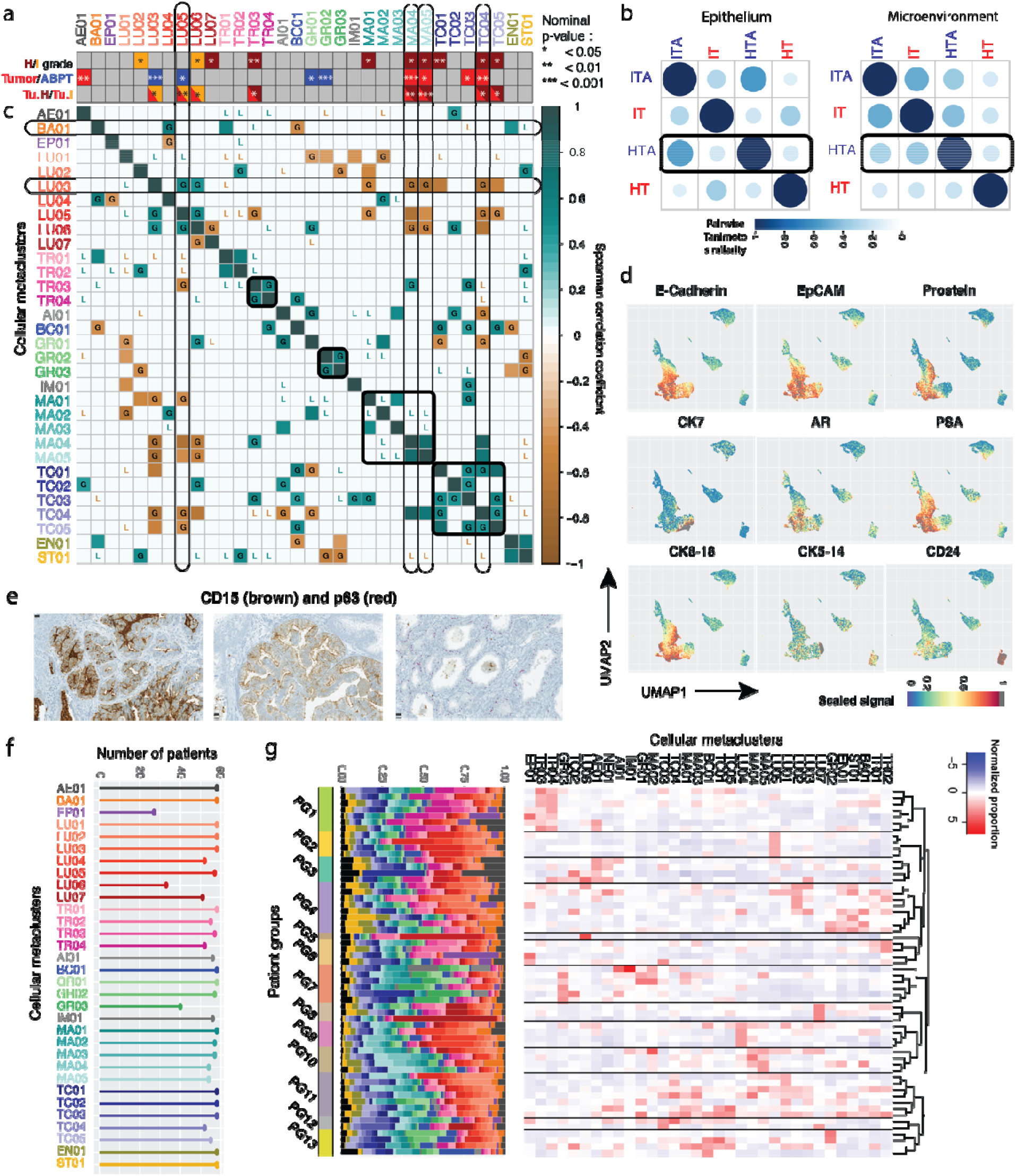
Characterization of epithelial tumor clusters and patient groups. (a) bar indicating which metaclusters were significantly enriched between three pairs of conditions: (top bar) high and intermediate grade samples, irrespective of tumor status; (middle bar) tumor versus ABPT; (bottom bar) intermediate and high grade tumor regions only. All nominal p values are given in Supplemental Table 2. (b) Pairwise Tanimoto similarity of intermediate (I) and high grade (H) tumor (T) and benign tumor-adjacent (TA) samples for metaclusters in the microenvironment and epithelium. (c) Correlation of metaclusters across 17 tumor patient samples. Correlations in the paired adjacent benign tissue that were lost in tumor are indicated by an L in the correlation plot while correlations that were gained are indicated by a G. Metacluster labels are colored to reflect cell types as in Figure 2a. (d) UMAP projections of 23200 cells (400 cells per patient) colored by expression of indicated epithelial and prostate-specific markers. Maximum signal (=1) is shown in grey. (e) CD15 and p63 co-stained showing CD15 expression in both a patient with acinar (left) and another with ductal (middle) carcinoma as well as absence of CD15 in normal glands (right) showing basal cell layer expressing p63. Scale 25μm. (f) Number of patients with cells belonging to a specific metacluster. Colors and labels matched to panels (a) and (c). (g) Grouped patient samples represented by proportion of metaclusters. Colors in bar plot reflect those on panels (a) and (c).

When interpreting changes in tumor and ABPT (Figure 4a, middle bar) in combination with tumor-only changes across grades (Figure 4a, bottom bar), we found that in some metaclusters (LU03, MA04, TC04) the effect between tumor and ABPT was stronger than the effect between patient grades. The depletion of LU03 and enrichment of MA04 and TC04 in tumors was observed across high and intermediate grade patients, however an effect could still be observed between the different grade groups. In other cases (AE01, GR01, GR02 and TC03), no effect could be detected between grades, only between tumor and ABPT samples, suggesting that enrichment (AE01 and TC03; p values = 0.003 and 0.022) or depletion (GR01 and GR02; p values = 0.044 and 0.0004) of these cellular phenotypes happens in tumors of patients irrespective of grade. Lastly, there were metaclusters that changed less significantly between tumor and ABPT (LU05, TR03 and MA05; p values = 0.025, 0.096 and 0.014) and more across grades (p values = 0.004, 0.026, and 0.0007). This suggests that although there may have been a difference between tumor and ABPT regions, a difference also existed between the tumor regions of intermediate and high grade tumors. These cellular phenotypes suggest a possible progressive change in the prostate, where some metaclusters are lowest in ABPT, higher in intermediate grade tumors, and even higher in high grade tumors (or the reverse with highest expression in ABPT and progressive loss in intermediate, then high grade tumors). We were also interested in integrating information across all cellular phenotypes in the epithelium and in the microenvironment (Figure 4b). We took the mean metacluster proportion across all 17 patients for which we had tumor and ABPT (tumor adjacent) samples. We calculated the Tanimoto similarity between intermediate tumor-adjacent regions (ITA), intermediate tumors (IT), high grade tumor-adjacent regions (HTA) and high grade tumors (HT). We found that in the epithelium, tumor-adjacent regions in intermediate and high grade are most similar to one another, while intermediate and high grade tumors bear more similarity to each other than to their benign adjacent regions. However, in the microenvironment, high grade tumor-adjacent regions (HTA) are more similar to intermediate grade tumors (IT) than to their paired high grade tumors (HT), suggesting that high grade tumors may have progressed from intermediate tumors, but with further changes in epithelial cellular phenotypes, while the tumor-adjacent microenvironment remains similar to that of an intermediate grade tumor (Figure 4b).

Next, we analyzed correlations between metaclusters in tumor samples across the 17 patients for which we had paired tumor and ABPT samples (Figure 4c). We restricted the analysis to Spearman correlations with a significance level < 0.05 for both correlations across tumor and ABPT samples. We compared correlations in the tumor to those in the paired adjacent benign tissue and found that while some correlations were lost in tumor (indicated by an L in the correlation plot; Figure 4c) others were gained (indicated by a G; Figure 4c). For example, luminal epithelial cell types LU03 and LU05 were uncorrelated in benign tissue but correlated in tumors. Indeed, we had found that both of these prostein-high metaclusters were depleted in tumors and we found their decrease was correlated with one another. We found that while transitional epithelial metaclusters TR01 and TR02 (CK7+/CK19+ cells and CK7+/CK19+ apoptotic cells respectively) were correlated in both tumor and adjacent benign prostate tissue, the strong correlation between TR03 and TR04 (CD15+/CK7+/CK19+ and CD15+/CK-low cells respectively) was only present in tumor regions. New correlations also appeared between cell types of the microenvironment. For example, GR02 and GR03 (CD15-low and CD3+ granulocytes, respectively) were newly correlated in tumor regions. We also observed that most T cell types became correlated in tumors although they were uncorrelated in adjacent benign tissue. Anti-correlations were also gained. Notably, MA04, MA05 (proliferating and CD45-low macrophages) and TC04 (proliferating T cells) became anti-correlated with LU03 in tumor. This result suggests that the increase of these macrophage and T cell metaclusters may be related to the depletion of this, likely benign, prostein-high luminal cell type.

While many new correlations and anti-correlations were gained in tumor regions, we also found that previously (anti-)correlated cell types in benign tissue became uncorrelated in tumor samples. Most macrophage metaclusters were correlated in benign tumor-adjacent samples but no longer in tumor. Basal cells were correlated with TR03 but this was not the case in tumor samples anymore, likely due to the de-regulated, malignant expansion of these transitional epithelial cells. Apoptotic epithelial cells (AE01) were correlated with transitional metaclusters TR01, TR02 and TR04 in tumor-adjacent regions but these correlations were lost in tumors. The main transitional epithelial metacluster TR01 used to be anti-correlated with various, likely benign, luminal cell types (LU01, LU03, LU05). It is believed that these transitional cells may originate from basal cells and are precursors of luminal cells and therefore a balance between the population of CK7+/CK19+ cells and CK8/18 exists in healthy scenario. However, our results support that this balance is disrupted in tumor regions.

The expression of CD15 in various malignant prostate epithelial cell populations (LU07, TR03 and TR04) was surprising although high-dimensional mapping using UMAP had already revealed CD15-high expressing cells in regions of epithelial marker expression such as E-Cadherin and EpCAM, luminal markers CK8 and CK18, and transitional epithelial markers CK7 and CK19 (Figures 3a and 4d). Immunohistochemistry confirmed the presence of CD15 in both ductal and acinar carcinoma of the prostate from patients who were shown to have this rare population by mass cytometry (one representative acinar and one ductal patient samples shown in Figure 4e). This adhesion molecule is typically used as a granulocyte marker and plays important roles in cell adhesion^16^ and migration^17^. Although previously observed in other carcinomas^18–20^ and demonstrated to be a marker of propagating tumor cells^21^, cells expressing CD15 had not been previously detected in prostate tumors. To assess the clinical relevance of this CD15+/CK19+ subpopulation (metacluster TR03) we analysed two other TMA cohorts with 374 patients (example core shown in <u>Supplemental Figure 7a</u>) with localized disease (336) and metastatic disease (38) and found that the proportion of patients with CD15 positive cells increased with disease severity (low, intermediate and high ISUP grades) and was highest amongst metastatic cases (Supplemental Figure 7b). Survival analysis did not yield any differences for this cohort however, the number of CD15+/CK19+ cases was very low. Only 5% positive cases were detected (19 patients out of 374, for which survival data was only available for 11) while via CyTOF most patients contained this cellular phenotype and 12% (7 out of 58) showed an enrichment. However, even in enriched cases, TR03 represented on average 0.6-1.1% of cells in a patient and overall, across all patients, this cellular phenotypes represented on average 0.3% of cells, making it difficult to identify a lot of potitive cases in a TMA spot with diameter0.06mm, here the number of cells is substantially lower than can be detected via high-throughput mass cytometry. It is very likely many CD15-positive patients were missed in the TMA analysis which impaired survival analysis.

Overall, 0.1%, 0.3% and 0.1% cells were found across the whole dataset from LU07, TR03 and TR04 respectively. Additionally, these rare CD15-expressing metaclusters were amongst the 14 observed in a subset of the patient cohort (Figure 4f). The remaining majority of metaclusters (29) were represented across all patients. In conclusion, our methodology identified a previously uncharacterized prostate-tumor subpopulation, which may also characterize a distinct patient subtype.

Although elevated PSA levels are typically associated with localized prostate cancer, luminal and transitional phenotypes found enriched in tumor or high grade samples had very little or no PSA expression (AE01, TR03 and LU07). Prostate cancer cells that express low levels or no PSA may be a self-renewing, tumor-propagating cell population that resists ADT^7^. Similarly, not all phenotypes increased in tumor regions were high in AR. While AR overexpression is associated with advanced disease and was found in one of the metaclusters enriched in high grade tumors (LU07), an AR-low phenotype was also enriched in high grade samples (TR03). Loss of AR has been associated with resistance to ADT^8^, and our results support that malignant phenotypes are not necessarily high in AR. We also found that AR-low metaclusters could be distinguished between benign and malignant by the presence of prostein and PSA. AR-low phenotypes that were PSA- and prostein-high were benign (LU03 and LU05), while malignant TR03, which was low in AR, was also low in PSA and prostein.

### Rare cellular phenotypes define patient subgroups

After having described the different cellular phenotypes present in prostate tumors, we wondered whether certain metaclusters (or combinations) could characterize patient groups. We clustered patients according to metacluster proportions using hierarchical clustering with Pearson correlation dissimilarity. We found many small groups across our 58 patient cohort that could each be characterized by the enrichment or depletion of a handful of phenotypes (Figure 4g). After statistical testing, we were able to define which metaclusters significantly defined a patient group (Figure 4g and Supplemental Figure 7c). Notably, patient group 1 consisted of patients with enrichment of transitional epithelial cells expressing CD15 (TR03 and TR04). The former, as had already been observed, was enriched in high grade tumors. Patient group 2 showed significant enrichment of prostein high phenotype LU05. Group 3 consisted of patients with the highest proportion of apoptotic epithelial cells (AE01), previously associated with tumor. One patient, with the highest enrichment of SMA, metacluster LU06, clustered separately from the rest and solely constituted patient group 5. High proportion of LU07 (CD15+, AR-high luminal cells), which we had already shown as enriched in high grade tumors (Figure 4a), was characteristic of patient group 8. The long-term effect that these phenotypes may have on survival remains to be determined. We showed that few single-cell phenotypes are necessary to further stratify patients beyond their ISUP tumor grade and may represent treatment targets for personalized treatment.

## DISCUSSION

We achieved the first single-cell analysis of prostate tumors and tissues by mass cytometry using a newly developed computational method that provides an unprecedented combination of high-dimensional clustering performance and speed. To reveal the phenotypic diversity of primary prostate tumors and their microenvironment, we profiled 1,670,117 cells from 58 prostate cancer patients through simultaneous quantification of 36 different protein abundances using mass cytometry.

For data analysis, we used the newly developed Franken pipeline. The initial step builds a SOM^22,23^ to over-cluster the preprocessed data into a large number of nodes. Next, a mutual *k*-nearest neighbor graph is created between the SOM nodes using the Tanimoto similarity^9^. This similarity measure is extremely suitable to analyzing high-dimensional data as it takes into consideration both the angle and the length of vectors when indicating their proximity making it more robust than more commonly used distance measures such as Euclidean or cosine. Lastly, the resulting graph is clustered using a random-walk-based graph clustering technique called Walktrap^10^. By comparing Franken to two state of the art methods for clustering mass cytometry data we found that when compared to FlowSOM, Franken provided superior F1-scores. Comparison of Franken to PhenoGraph showed that although they performed equivalently in F1-scoring for data-sets of around 200,000 cells, Franken could be run on up to 40 million cells (which would be computationally infeasible for PhenoGraph) in the time it would take Phenograph to analyze 1 million cells. Franken also ran over 20 times faster than the state-of-the-art single-cell RNA sequencing clustering technique, Seurat. Our Franken pipeline was able to identify very rare subpopulations with a frequency as low as 1/5,000 and detected previously unknown rare prostate cancer phenotypes which were later confirmed through imaging, showing that Franken does not compromise performance or sensitivity while providing high scalability.

As Franken is sensitive, it can detect rare metaclusters based on subtle gradient variation of markers. To focus on clusters with qualitatively similar expression patterns through hierarchical clustering of correlation similarities and described 33 prostate cellular phenotypes including 14 epithelial and 18 cell types from the microenvironment and one cluster with very low to no expression of the markers in our panel. Tumors and surrounding ABPT had considerable similarity: Nine of the 33 metaclusters (27%) were present at significantly different frequencies between the two regions. Although this was much higher than would be expected at random for a confidence level of 5% (1.65 out of 33), almost two thirds or cell types and states detected (24 out of 33), were shared at similar proportions in tumor and ABPT tissue. Most epithelial differences between tumor and ABPT and tumor grades involved luminal cell types with the exception of one transitional epithelial phenotype (the CD15-high metacluster TR03), suggesting that prostate tumorigenesis is strongly affected by an interplay of luminal phenotypes. Prostein-high phenotypes were depleted in tumor regions and even more so in high grade tumors; this suggests that during tumorigenesis there is selection for poorly differentiated cell types. Tumor-enriched phenotypes all contained EpCAM. High levels of EpCAM expression at both mRNA and protein levels were previously reported in prostate cancer tissues and cell lines^24,25^. Our analysis showed that both AR-high /PSA-low (LU07) and AR-low /PSA-low (TR03) cells were present in localized, hormone-naïve prostate tumors even though such phenotypes had previously been associated only with castration-resistant disease after ADT or metastatic disease ^6–8^. Cells in the AR-high /PSA-low cluster also overexpressed Nkx3.1. Loss of prostein^26^ and PSA expression^27^, Nkx3.1 overexpression^28^, and AR overexpression or amplification^29–31^ have been shown to be common in castration-resistant disease states. Furthermore, men with localized high-grade prostate cancer but low PSA show inferior cancer survival^32^. It remains to be determined whether these rare cells with the properties of tumors resistant to ADT are capable of dissemination and are responsible for disease progression after prostatectomy. Surprisingly, two phenotypes enriched in high grade patients expressed CD15. After analysing an additional 374 patients’ TMA samples, we also found that CD15+/CK19+ prostate epithelial cells were further enriched in metastatic disease. CD15 plays an important role in cell adhesion and migration^16,17^ and CD15-expressing cells have been identified in other tumor types as having stem-like potential but not in prostate cancer^18–20^ and might represent a new biomarker for aggressive phenotypes with a bigger potential to metastasize.

The tumor microenvironments were similar in both the tumor regions and the neighboring ABPT regions for patients of different tumor grades with the exception of granulocytes, which were present at lower levels in tumors regardless of grade. Other rarer immune cell types changed both between tumor and ABPT regions as well as across tumor grades. In particular, we observed that one proliferating T-cell (TC04) and two macrophage (proliferating MA04 and CD45-low MA05) phenotypes were enriched in tumor regions and were further enriched in high-grade tumors.

The microenvironments of tumors from the intermediate sub-cohort had lower frequencies of immune phenotypes compared to the ABPT regions, but we found the opposite in the high grade sub-cohort. In high grade tumors, there was an enrichment of multiple immune phenotypes compared to the ABPT regions. It is currently unclear whether intermediate grade tumors progress to high grade disease. If such a progression happens, our analysis suggests that the hyperplasia and expansion of the epithelial compartment might precede alterations in the tumor microenvironment, or it may be that these differences are reflective of disease stage. We proceeded to analyze the overall changes in the microenvironment and epithelium by integrating information across all metaclusters in these two compartments and estimating the Tanimoto similarity between high grade (HT) and intermediate tumors (IT) as well as high grade and intermediate tumor-adjacent ABPT regions (HTA and ITA, respectively). We found further evidence of a possible progression from intermediate to high grade tumors suggested by the similarity of the microenvironments of intermediate grade tumors and high-grade ABPT regions. Some of the rare cell types in the microenvironment, which we found to be enriched in tumors from both grade groups and further enriched in high grade patients may represent new putative targets that can be used to prevent the progression of the disease.

Tumors do not grow in isolation; cancerous cells require support from the microenvironment. Accessory cells have been successfully targeted with therapy^33–35^. We found that most immune metaclusters were present at similar frequencies or were decreased in the tumor compared to the adjacent ABPT regions with the notable exceptions of rare T cell and macrophages cellular phenotypes. Both monocyte infiltration and macrophage proliferation are necessary for macrophage maintenance during tumor growth^36^, and in breast cancer, proliferating macrophages are associated with high tumor grade, hormone receptor negativity, and poor clinical outcome^37^. However, macrophage counting based on immunohistochemical analysis had not led to any consensus on the prognostic significance of tumor-associated macrophages in ^12,38^ prostate cancer. In our prostate cancer cohort, proliferative macrophages were enriched in prostate tumors and even more so in high grade patients. Taken together with findings that tumor-associated macrophages (TAMs) are capable of proliferation^37^, our data suggests that in addition to therapy that inhibits differentiation of TAMs from circulating monocytes, blocking the proliferation of macrophages may have an effect in slowing the development of high grade disease in prostate cancer. Our clustering analysis identified that not all macrophage phenotypes changed frequency in tumor compared to ABPT regions. This suggests the presence of separate cancer-versus stroma-infiltrating macrophage phenotypes that may have opposing influences on tumorigenesis^12,14^, highlighting the importance of investigating macrophage infiltration in prostate cancer.

In summary, new biomarkers are needed to identify which men qualify for active surveillance or need aggressive treatment. Understanding the cellular complexity of prostate tumors and their microenvironments is key to the development of new diagnostic and treatment strategies. Here, we provide a thorough description of prostate tissue heterogeneity on the single-cell level and describe differences between tumors and the neighboring benign hyperplasia regions as well as across patient grades. We identify two CD15-high phenotypes enriched in high grade patients as well as changes to the microenvironment in rare macrophage and T cell phenotypes associated with tumor regions and high grade disease. We also identify in men with localized disease, epithelial subpopulations associated with advanced castration-resistant disease. The alterations to the epithelium and microenvironment should be further explored to guide development of new diagnostic and treatment paradigms for prostate cancer and to understand which cellular phenotypes in primary prostate cancer need to be detected and may change treatment decisions.

## METHODS

### Patient samples and tissue microarray construction

The Ethics Committee of the Canton of Zurich approved all procedures involving human prostate material (KEK-ZH-No. 2008-0040). All patients were part of the Zurich Prostate Cancer Outcomes Cohort (ProCOC) study ^39,40^, and each patient signed an informed consent form. Prostatectomy samples were taken from 58 prostate cancer patients from the ProCOC cohort between 2015 and 2017. Tumors were of a range of ISUP grades. No clinical or histological status was used in the selection of the cohort. Staging and grading was performed using World Health Organization and ISUP criteria^41^). Twenty-four patients had ISUP grade II (Gleason score 3+4), 22 had ISUP grade III (Gleason score 4+3), and 12 had ISUP grade V prostate carcinoma (Gleason scores 4+5, 5+4, and 5+5).

Immediately after surgery, native radical prostatectomy specimens were transferred to the frozen section lab on ice (4 °C) and were processed within 15 min in the Department of Pathology and Molecular Pathology, University Hospital Zurich. The first slice after dissection of the apex was quartered and snap frozen in four separate blocks for biobanking within the ProCOC study. Fresh tumor and ABPT tissue was taken from the second slice after dissection of the apex without destruction of surgical margins and the pseudocapsule. After formalin fixation overnight, the rest of the specimen was embedded in paraffin. Hematoxylin and eosin stained sections of the four frozen blocks were sliced for immediate evaluation regarding tumor load and margins in synopsis with the standard formalin-fixed paraffin-embedded histology to control for the representativeness of tissue sampling for mass cytometry.

Following evaluation of tissue sections by uropathologists (NJR, JHR, PJW) a tissue microarray (TMA) containing two ABPT and two tumor regions from all patients in the selected cohort was generated as previously described ^42^. For TMA construction, representative tumor areas of the second and third slice of radical prostatectomy specimens were chosen, as close as possible to the area of tissue sampling for mass cytometry. Supplementary Figure 5 shows H&E images of the selected regions.

### Fresh tissue preparation

After surgical resection and based on the aforementioned real-time frozen sections, the index tumor lesions (the most extensive with the highest Gleason score) were immediately harvested and transferred to precooled MACS tissue storage solution (Miltenyi Biotec) and shipped at 4°C. To better select the index lesion, only cases in which a tumor nodule was also macroscopically visible were selected, ultimately resulting in a cohort with higher ISUP grades. Tissue processing was completed within 24 h of collection. For the dissociation of tissues to single cells, the tissue was minced using surgical scalpels and further disintegrated using the Tumor Dissociation Kit, human (Miltenyi Biotech) and the gentleMACS Dissociator (Miltenyi Biotech) according to the manufacturer’s instructions. The resulting single-cell suspensions were filtered through sterile 70-μm and 40-μm cell strainers and stained for viability with 25 μM cisplatin (Enzo Life Sciences) in a 1-min pulse before quenching with 10% FBS (Fienberg et al., 2012). Cells were then fixed with 1.6% paraformaldehyde (Electron Microscopy Sciences) for 10 min at room temperature and stored at −80 °C.

### Mass cytometry barcoding

To ensure homogenous staining, 0.3 × 10^6^ to 0.8 × 10^6^ cells from each tumor sample were barcoded as previously described using a 126-well barcoding scheme consisting of unique combinations of four out of nine barcoding reagents^43^. Metals included palladium (^105^Pd, ^106^Pd, ^108^Pd, ^110^Pd, Fluidigm) conjugated to bromoacetamidobenzyl-EDTA (Dojindo) and indium (^113^In and ^115^In, Fluidigm), yttrium (^89^Y, Sigma Aldrich), rhodium (^103^Rh, Sigma Aldrich), and bismuth (^209^Bi, Sigma Aldrich) conjugated to maleimido-mono-amide-DOTA (Macrocyclics). The concentrations were adjusted to 20 nM (^209^Bi), 100 nM (^105-110^Pd, ^115^In, ^89^Y), 200 nM (^113^In), or 2 μM (^103^Rh) as previously reported to be optimal^44^. Cells were barcoded using the transient partial permeabilization protocol^45^. Cells were washed with 0.03% saponin in PBS (Sigma Aldrich) and incubated for 30 min at room temperature with 200 μl of mass tag barcoding reagents. Cells were then washed twice with PBS plus saponin and twice with cell staining medium (CSM, PBS with 0.5% bovine serum albumin and 0.02% sodium azide).

### Antibodies and antibody labeling

The supplier, clone, and metal tag for each antibody used in this study are listed in Supplemental Table 3. Antibody labeling with the indicated metal tag was performed using the MaxPAR antibody conjugation kit (Fluidigm). After metal conjugation, the concentration of each antibody was assessed using a Nanodrop (Thermo Scientific). The concentration was adjusted to 200 μg/ml and stored in Candor Antibody Stabilizer. All conjugated antibodies were titrated for optimal concentration for use with prostate tissues. Antibody usage in this study was managed using the AirLab cloud-based platform^46^.

### Immunohistochemistry

For immunohistochemical validation studies anti-CD3 (mouse monoclonal, clone LN10, Leica Microsystems) and anti-CD15 (mouse monoclonal, clone Carb-3, Agilent Dako) antibodies were used. Automated platforms were used for in situ protein expression analyses of CD15 (Ventana Benchmark CD15), and CD3 (Leica Bond-Max).

### Antibody staining and mass cytometry data collection

After barcoding, pooled cells were incubated with FcR blocking reagent (Miltenyi Biotech) for 10 min at 4 °C. Samples were stained with 100 μl of the antibody panel per 10^6^ cells for 60 min at 4 °C. Cells were washed twice in CSM and resuspended in 1 ml of nucleic acid Ir-Intercalator (Fluidigm) overnight at 4 °C. Cells were then washed once in CSM, once in PBS, and twice in water. Cells were then diluted to 0.5 × 10^6^ cells/ml in H_2_O containing 10% of EQ™ Four Element Calibration Beads (Fluidigm). Samples were placed on ice until analysis. Data were acquired on an upgraded Helios CyTOF 2 mass cytometer using the Super Sampler (Victorian Airship) introduction system.

### Mass cytometry data analysis

Individual .fcs files collected from each set of samples were concatenated using the .fcs concatenation tool from Cytobank, and data were normalized using the executable MATLAB version of the Normalizer tool ^47^ Individual samples were debarcoded using the CATALYST R/Bioconductor package ^48^. Debarcoded files were compensated for channel crosstalk using single-stained polystyrene beads as previously described ^48^.

CyTOF data was analyzed by initially applying an arcsinh transformation with a cofactor of 5 *(newdata_i_ = arcsinh(data_i_/5)).* The UMAP algorithm^49^ was applied to the high-dimensional data from 23,200 (400 per patient) cells taken at random from across the patient cohort using default parameters (perplexity, 30; theta, 0.5) to facilitate visualization in two dimensions. The pre-processed data were analyzed using the Franken algorithm as described in detail below. All analysis was done using R version 3.4.1

### The Franken pipeline

The initial step of the Franken pipeline uses a SOM ^22,23^ to over-cluster the preprocessed data into a large number of nodes. Prostate patient data was pooled from all patient samples (1,670,117 cells) and 400 SOM nodes were used (SOM grid dimensions were x=20 and y=20). Next, a mutual *k*-nearest neighbor graph (k=6) was built between the SOM nodes using the Tanimoto similarity ^9^, which unlike the binary version, can be applied to continuous or discrete non-negative features and retains the sparsity property of the cosine while allowing discrimination of collinear vectors. Lastly, the resulting graph is clustered using a random-walk-based graph clustering technique called Walktrap, using the implementations available in package igraph ^50^ in R. Walktrap is a graph partitioning technique that requires a choice of random walk steps. To increase our pipeline’s robustness, this procedure is applied for a range of random walk steps and the smallest step that maximizes the graph’s modularity is chosen.

According to the thorough review and comparison of community detection algorithms by Yang et al. ^51^, Walktrap is amongst the best performing algorithms for both large and small networks regardless of whether the mixing parameter is high or low. Although Yang et al. found that for large mixing parameters most algorithms failed to detect the community structure, Walktrap was able to do it. Another advantage of Walktrap is that it is possible (although not necessary) for a user to define the number of communities one wishes to find in the data. This allows the user to decide exactly how many clusters they wish to find, although the method is by default run in an unsupervised way. Although Walktrap is not the fastest method for large networks, the network size in Franken is never large due to the initial SOM-building step.

### Benchmarking Franken against other methods for additional datasets

To test the performance of Franken, we compared it to two state-of-the-art clustering methods for mass cytometry data, Phenograph^52^ and FlowSOM^23^. All three methods were used to cluster data obtained from analysis of healthy bone marrow cells^53^ and data from 10 cell lines stained with our prostate-centric antibody panel (Supplemental Figure 1b and 1c). The cell phenotypes were manually annotated in the data obtained from analysis of the bone marrow cells according to Bendall et al.^53^. We calculated precision and recall (as shown by F1 scores according to Weber et al.^5^) for each phenotype in each dataset (Supplemental Figure 1a and 1b). Franken was able to recover the most phenotypes in both datasets, resulting in the least phenotypes with zero F1 scores. After repeating the F1 estimates for multiple runs (with different random seeds) of each method, on average, Franken performed as well or better than the other methods. Franken requires the input of three parameters: the SOM grid dimensions (x and y which multiplied correspond to the number of nodes used to build the SOM) and k neighbors (the number of neighbors used to decide whether two nodes are connected by an edge in the mutual nearest neighbor graph); a SOM size of 400 (x=y=20) and k = 6 were used in our simulations of all three datasets. Franken results were robust to the choice of its parameters (Supplemental Figure 1).

Franken requires minimal computational resources, and runtimes were very fast when evaluated on the two benchmark datasets containing around 200,000 cells (Supplemental Figure 2d). While FlowSOM was slightly faster than Franken, it performed very poorly in F1 scores compared to both Franken and Phenograph. FlowSOM was run using the default parameters chosen by the authors as optimal (SOM nodes 100). However, as FlowSOM and Franken share the SOM building step we also tested FlowSOM using the same 400 nodes SOM grid however the F1 scores were equivalent and the runtime was increased, no longer making FlwoSOM superior in speed therefore we chose to use the author’s default FlowSOM parameters. We hypothesize that the poor performance of FlowSOM is due to the hierarchical clustering step which is less suitable to the high dimensional node representation resulting from the SOM than our mutual nearest neighbor graph-building approach.

Although the F1 scores from Phenograph were comparable to Franken’s, their scalability varied greatly. After testing Franken on several synthetic datasets of sizes varying from 20 thousand to 40 million, we showed that one could analyze 40 million cells with Franken in the equivalent time taken to analyze 1 million cells using Phenograph (Supplemental Figure 2). Franken can also be applied to single-cell RNA sequencing data, therefore we also compared Franken’s scalability with the state-of-the-art method for single-cell RNA sequencing Seurat and could show that Franken was far superior in scalability (Supplemental Figure 2). Franken could cluster 40 million cells in the half of the time taken to cluster 3 million cells with Seurat. As Phenograph and Seurat require far larger computational resources they could not be run on the larger datasets beyond 1 and 3 million respectively.

### Other computational methods

PhenoGraph runs included in Supplemental Figure 1 were performed using the MATLAB (R2018b) implementation using the GUI CYT3 as the matlab implementation was the only one which allowed different random seeds to be used in each run. Default parameters were used: k nearest neighbors = 30. PhenoGraph runs in supplementary Figure 2 were performed using its implementation in R (Rphenograph) for its ease in including it in scripts instead of manually running the MATLAB GUI.

FlowSOM and Seurat runs were performed using their implementation in R and default parameters.

### TMA analysis

Immunohistochemistry applied to TMA was used to validate single-cell mass cytometry data. The open-source software QuPath ^54^ was used to quantify cell types in TMA. CD3^+^ cells were quantified using an automated detection procedure, and CD15^+^ cells were manually selected by a pathologist (J.H.R.).

## Supporting information

ST1 cell lines

ST2 pvals

ST3 antibodies

## Data availability

The single-cell data supporting the findings of this study including raw .fcs files from primary samples and cell lines as well as TMA images will be available online upon publication.

The Bone marrow CyTOF data pertaining to Figure 2 refers to article: S.C. Bendall, E.F. Simonds, P. Qiu, A.D. Amir, P.O. Krutzik, R. Finck, R.V. Bruggner, R.Mela med, A. Trejo, O.I. Ornatsky, *et al.* Single-cell mass cytometry of differential immune and drug responses across a human hematopoietic continuum Science, 332 (2011), pp. 687-696. The raw data is publicly available at http://cytobank.org/nolanlab/reports and was pre-processed according to Weber, Lukas M; Robinson, Mark D (2016). Comparison of clustering methods for high-dimensional single-cell flow and mass cytometry data. Cytometry Part A, 89(12):1084-1096.

## Code availability

Code for data pre-processing pipeline (from Weber et al.) can be found here: https://github.com/lmweber/cytometry-clustering-comparison

The Franken package is available at https://github.com/ldvroditi/Franken

## ACKNOWLEDGEMENTS

The research was supported by a European Research Council PrECISE project from the European Union Horizon 2020 research and innovation program under grant agreement No. 668858. We thank the Wild and Bodenmiller lab members as well as Dr. Andreas Moor for useful discussions and their valuable feedback.

## AUTHOR CONTRIBUTIONS

L.D.V.R., B.B., and P.W. conceived the study. P.W., B.B., A.J. and L.D.V.R designed the antibody panel and A.J. performed all antibody validation and data acquisition experiments. L.D.V.R. designed and developed Franken and performed data analysis. L.D.V.R and S.C tested algorithm in multiple datasets. T.H., C.P. and C.D.F. provided clinical samples. L.D.V.R. and P.W. developed TMA. P.W. P.B, J.H.R., A.T. and L.D.V.R. performed the TMA IHC image analysis. L.D.V.R. B.B. and P.W. performed biological analysis and interpretation with input from J.H.R and H.W.J.. L.D.V.R, B.B. and P.W. wrote the manuscript with input from F.C., H. W.J, S.C and C.D.F.

## COMPETING INTERESTS

The authors declare no competing interests.

## Notes

http://cytobank.org/nolanlab/reports

https://github.com/lmweber/cytometry-clustering-comparison

https://github.com/ldvroditi/Franken

### Competing Interest Statement

The authors have declared no competing interest.

