## Supplement legends for "Single-Cell Proteomics Defines the Cellular Heterogeneity of Localized Prostate Cancer"

**Figure S1.** **High-dimensional clustering with Franken combines performance and speed.** (a) Franken involves fitting a large SOM to the data, building a mutual *k*-nearest neighbor graph on the *n*-dimensional SOM nodes, and applying the Walktrap algorithm. A UMAP 2-D map can be used for visualization. (b-c) Performance of Franken, PhenoGraph, and FlowSOM in clustering of b) bone marrow and c) cell lines as measured by F1 scores. (d) Average runtimes of Franken, PhenoGraph, and FlowSOM applied to the bone marrow (blue) and cell line (pink) datasets 10 times and plotted against mean F1 score. FlowSOM and PhenoGraph were run using their default parameters in their R and Matlab implementations, respectively (e) F1 values for Franken tested on cell lines dataset for different SOM sizes as well as number of k neighbors and distance measures in mutual nearest neighbor graph. The extended Jaccard measure (ejaccard) is the recommended measure for Franken. The default k-nearest neighbors is kn=6. As Franken builds a mutual k-nearest-neighbor graph, too few neighbors can lead to poor results, however Franken proved to be very stable a minimum number of neighbors of at least kn = 3. (f) Franken, PhenoGraph and FlowSOM were run on a series of synthetic datasets of increasing size up to 40 million cells and runtimes were recorded for each. These consisted of simulated gaussians in 10 dimensions. We also included a state-of-the-art clustering technique used for single-cell RNA sequencing data (Seurat) according to Duo et al. 2018. PhenoGraph and Seurat could not be run on the largest datasets and were therefore run on the largest computationally feasible set (1 million for PhenoGraph and 3 million for Seurat). Franken was able to cluster 40 million cells in the same time necessary for PhenoGraph to cluster 1 million cells. Seurat performed slightly faster than PhenoGraphbut attempting to cluster 3 million cells with Seurat took twice as long as it would take to cluster 40 million cells with Franken. FlowSOM was faster than all methods, however its F1-score performance was far inferior compared to Franken and PhenoGraph when applied to all benchmarking datasets (Figure 2b-d). Franken still provided comparable speed to FlowSOM and was at most three times slower for the largest 40 million dataset.

**Figure S2. Logistic regression classification confirms metaclustering resolution.** Metacluster labels given in b-e are all given in panel (d). (a) UMAP 2-D representation colored by only epithelial, immune or microenvironemnt cell subsets (b) Number of cells in each metacluster (c) Proportion of metacluster (d) Misclassification error for each metacluster resulting from logistic regression classification. (e) Coefficients from logistic regression performed with LASSO regularization.

**Figure S3. Prostate patients share a lot of metaclusters.** (a) Relative proportion of cells in a metacluster from a tumor-sample (red), adjacent bening prostatic tissue-sample (ABPT; blue) and from a random prostatic tissue (RPT; grey) (b) Average proportion (normalized by total number of cell in a patient) of cell types across all 58 patients in cohort (c) Expression of Stromal (SMA, S100A4 and Vimentin) and endothelial (CD31) markers (d) H&E of TMA from patient cohort analysed with CyTOF. The TMA contains two BPH and two tumor regions from all patients in the selected cohort was generated as previously described (Mortezavi et al. 2011). For TMA construction, representative tumor areas of the second and third slice of radical prostatectomy specimens were chosen, as close as possible to the area of tissue sampling for mass cytometry.

**Figure S4. Immune metacluster comparison between grades and tumor/non-tumor** **groups**. Proportions (normalized by total number of cells in a patient) of (a) T cell, (b) macrophages and (c) granulocytes metaclusters across all 58 patients in cohort (Intermediate N = 46 and high grade N = 12) (d) Proportion (normalized by total number of cells in a patient) of cells from tumor and adjacent benign prostatic tissue samples for each granulocyte metacluster (N=17). Dots are colored by disease severity (intermediate vs high grade). Paired tumor/ABPT samples were analysed with a two-sided Wilcoxon signed rank paired test and unpaired intermediate/high grade samples were analysed with a two-sided Wilcoxon rank sum test (also known as a Mann-Whitney-Wilcoxon).

**Figure S5. Epithelial metacluster comparison between grades and tumor/non-tumor** **groups**. (a) Comparison of proportions (b) Proportion (normalized by total number of cells in a patient) of luminal cells metaclusters stratified by (c) tumor and adjacent benign prostatic tissue samples (N=17) and (d) intermediate and high grade patient samples (for combined tumor/ABPT; Intermediate N = 46 and high grade N = 12). (e) same as (d) for transitional epithelial metaclusters. (e) roportion (normalized by total number of cells in a patient) of cells from tumor and adjacent benign prostatic tissue samples for each transitional epithelial metacluster. Dots are colored by disease severity (intermediate vs high grade). Paired tumor/ABPT samples were analysed with a two-sided Wilcoxon signed rank paired test and unpaired intermediate/high grade samples were analysed with a two-sided Wilcoxon rank sum test (also known as a Mann-Whitney-Wilcoxon).

**Figure S6. UMAP 2-D representation of cells across all 58 patients.** 400 cells were sampled from each patient and projected using dimensionality reduction with UMAP. The normalized expression of each marker is shown for each cell.

**Figure S7. Analysis of CD15 prostate epithelial cells and patient groups.** (a) Sample core from TMA stained with CD15 (brown) and CK19 (red). The diameter of each spot is exactly 0.6 mm (b) The TMA was graded as positive or negative in the presence or absence (respectively) of double CD15 and CK19 positivity. The bar plot represents the proportion of patients positive for the double staining. From low grade (Gleason 6 or lower) to metastatic patients the proportions are (Positive/Negative): 1/47 (low grade); 11/188 (intermediate grade); 7/88 (high grade) and 4/34 (metastatisis). (c) Summary of significance testing showing which cellular metaclusters are enriched or depleted in a patient group. Only significant p-values below of at least 0.05 are shown.
