## Supplementary figures and images for "Single-Cell Proteomics Defines the Cellular Heterogeneity of Localized Prostate Cancer"

### Fig_S1

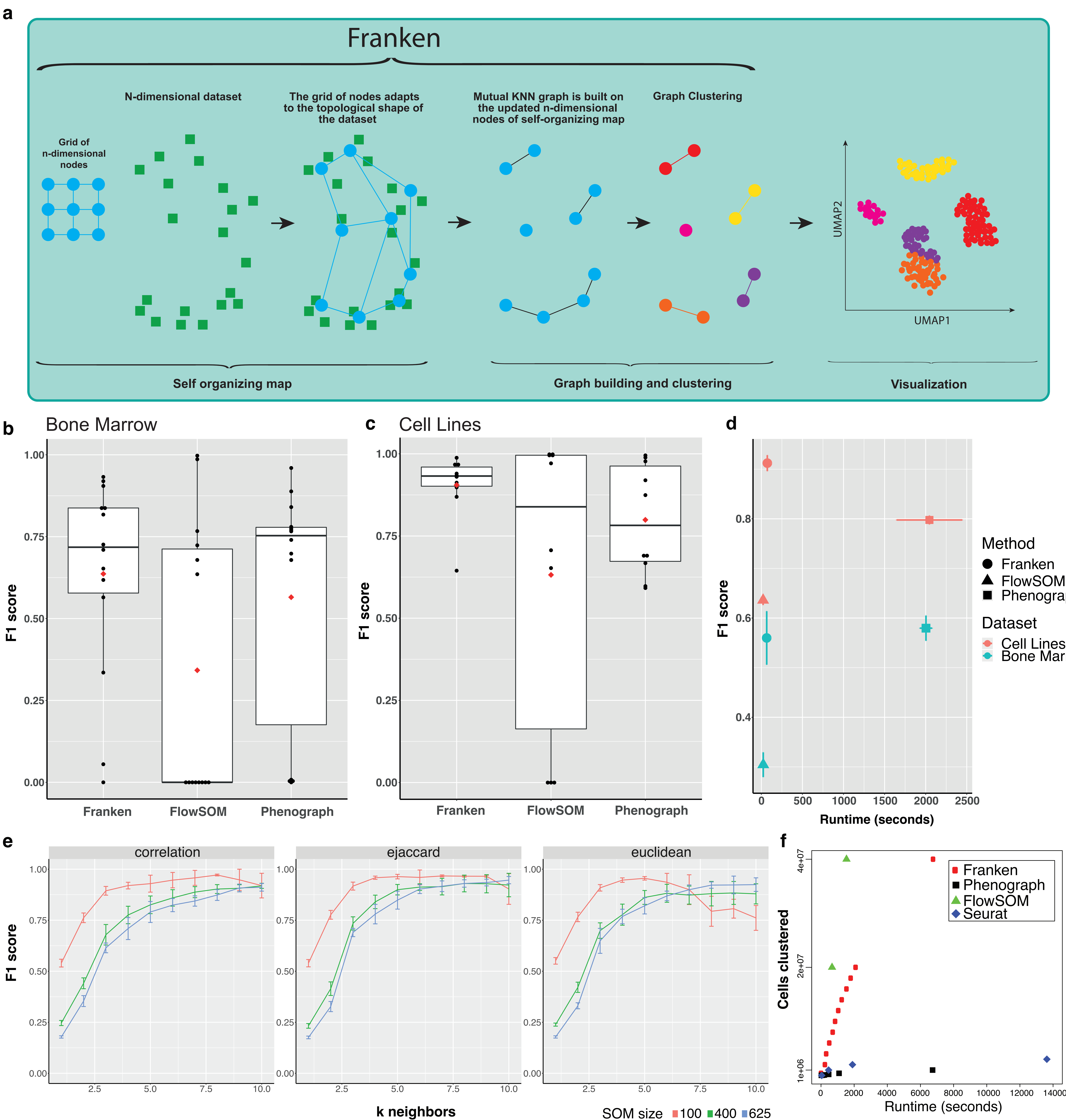

### Fig_S2

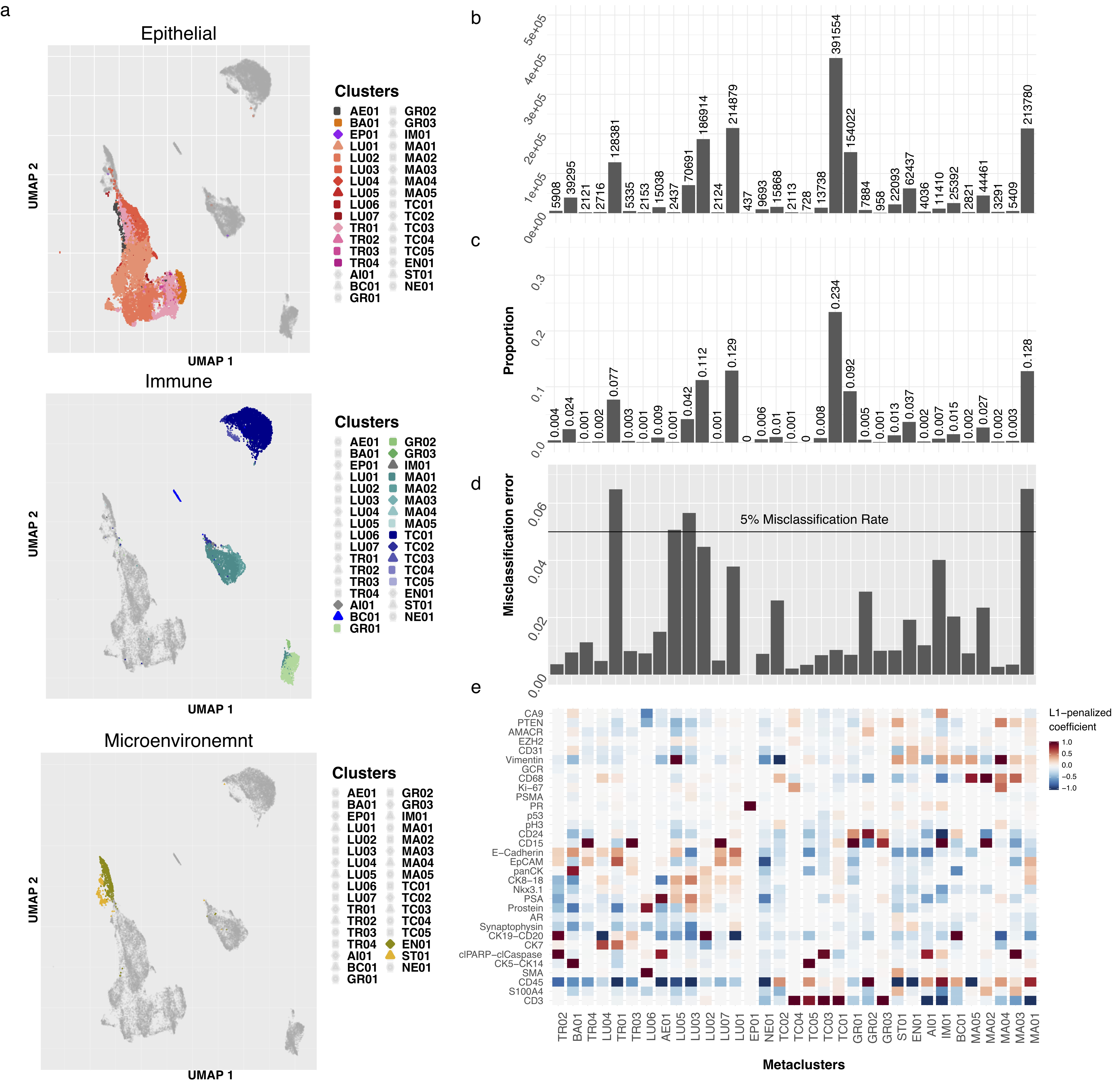

### Fig_S3

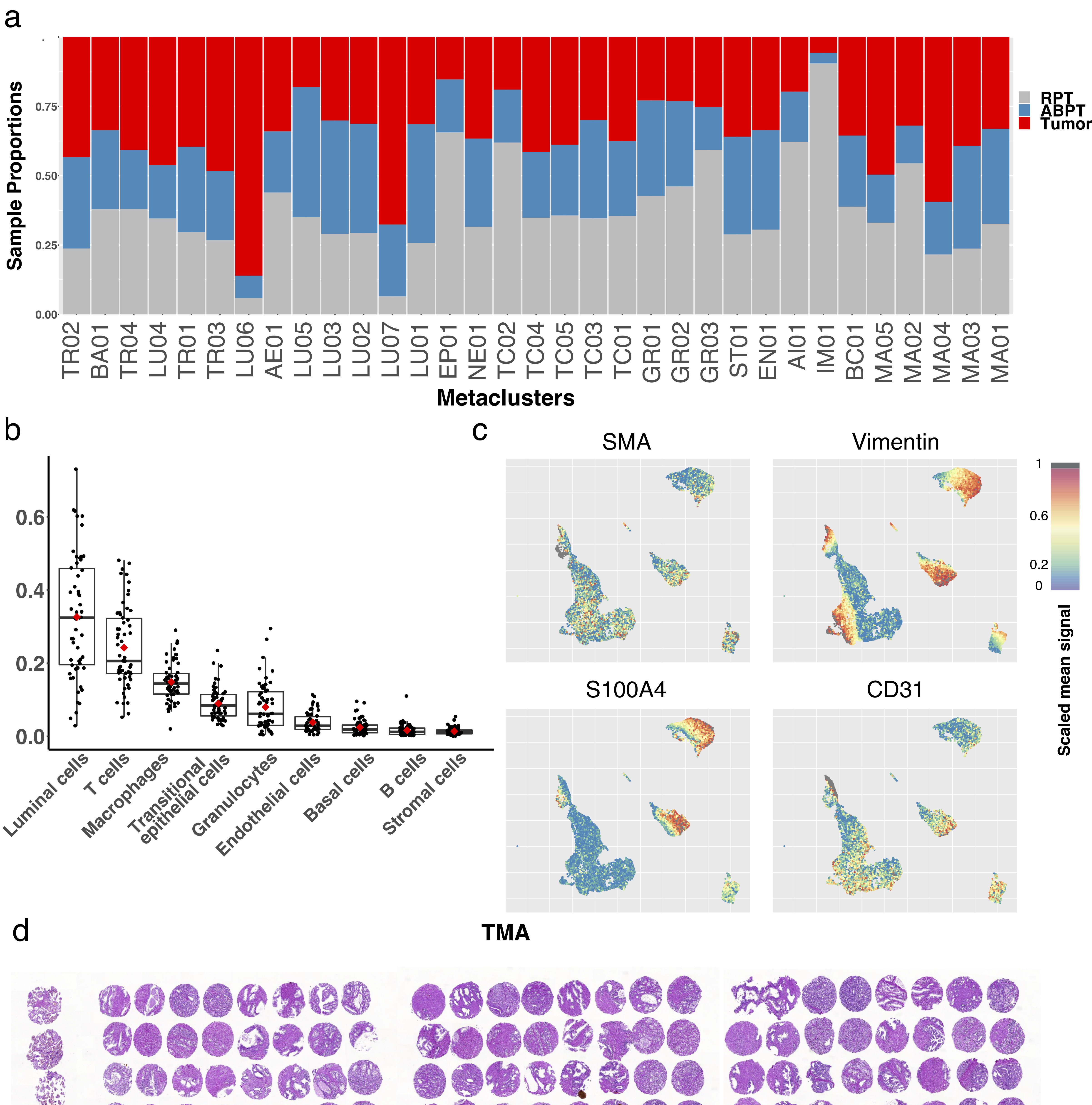

### Fig_S4

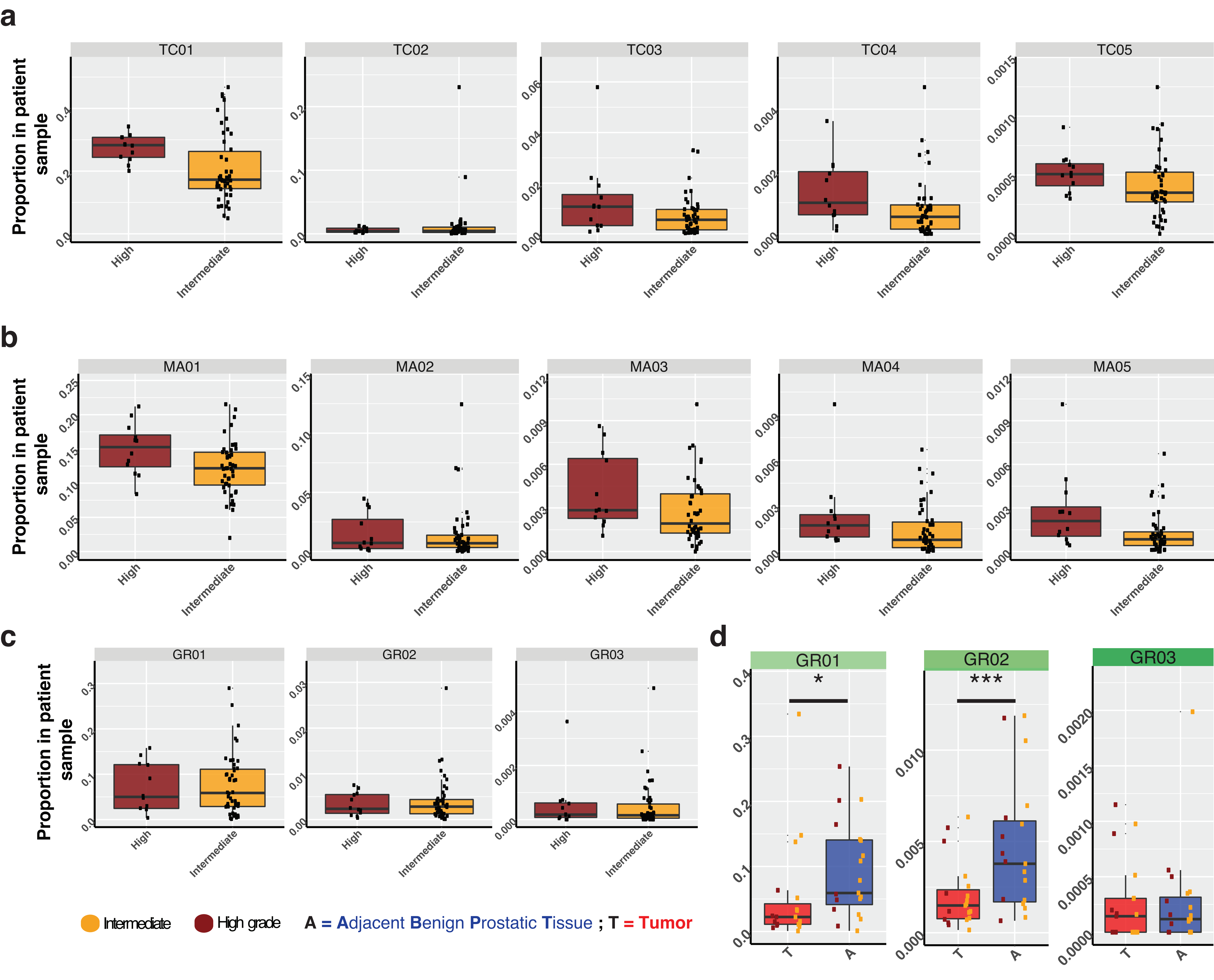

### Fig_S5

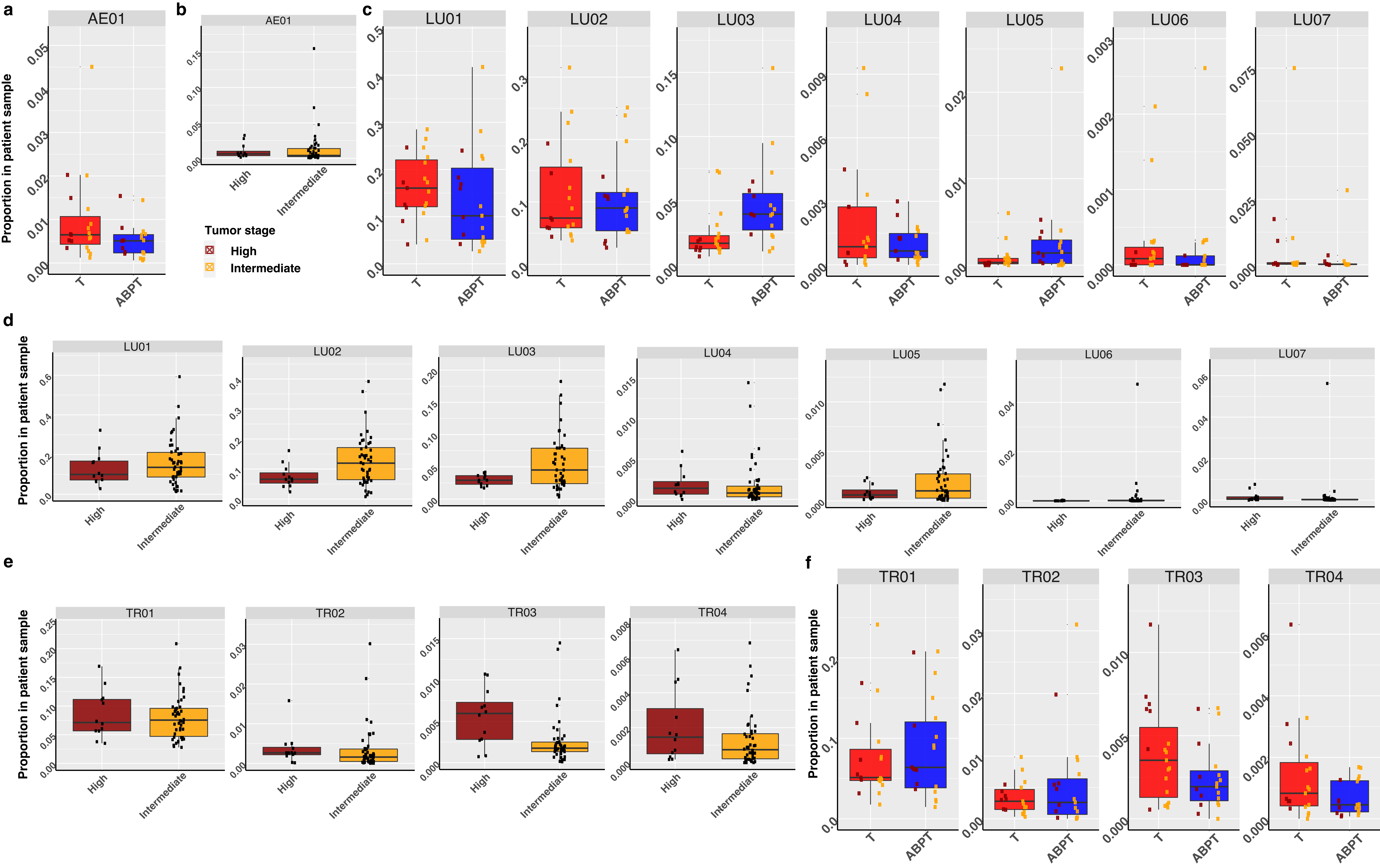

### Fig_S6

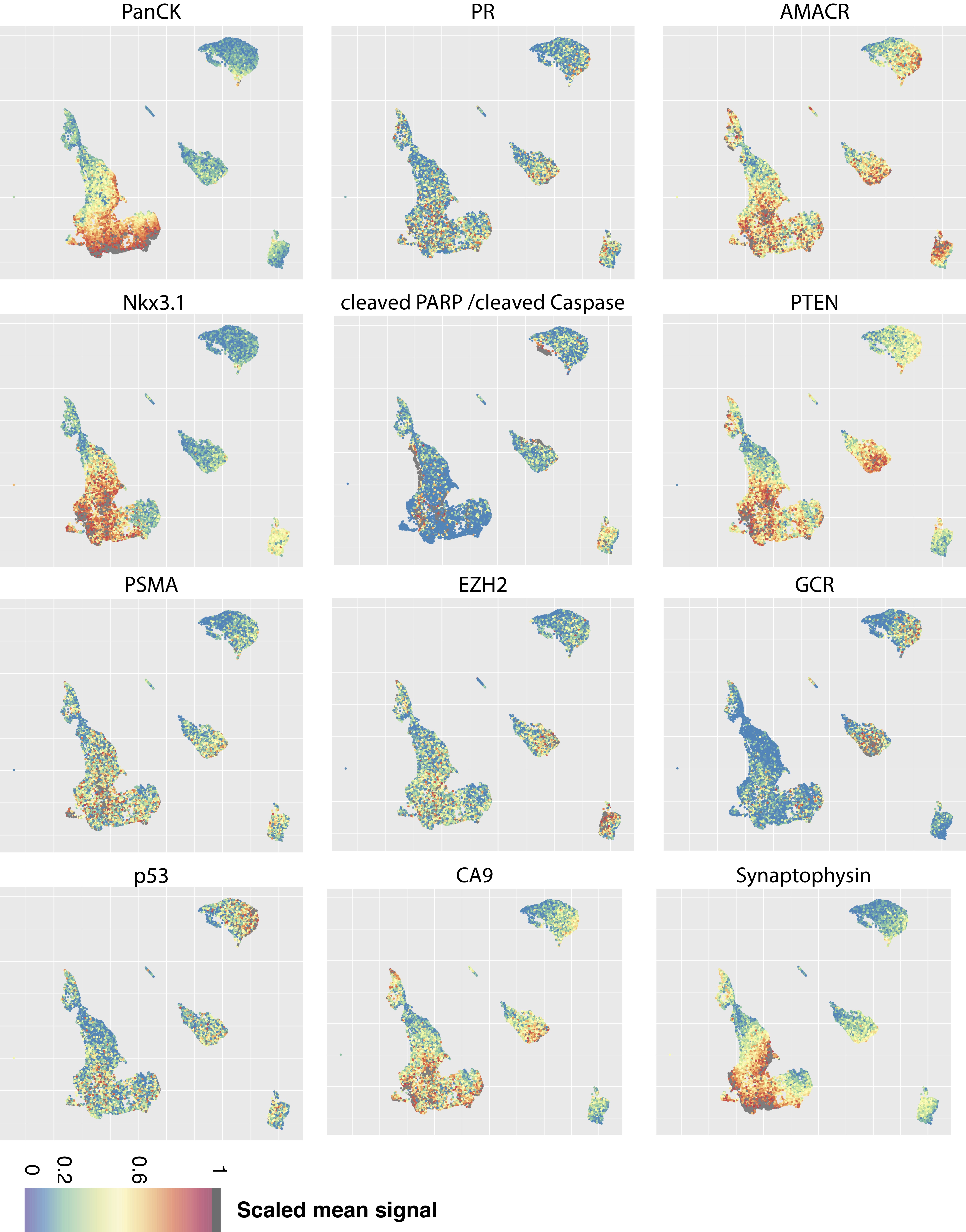

### Fig_S7

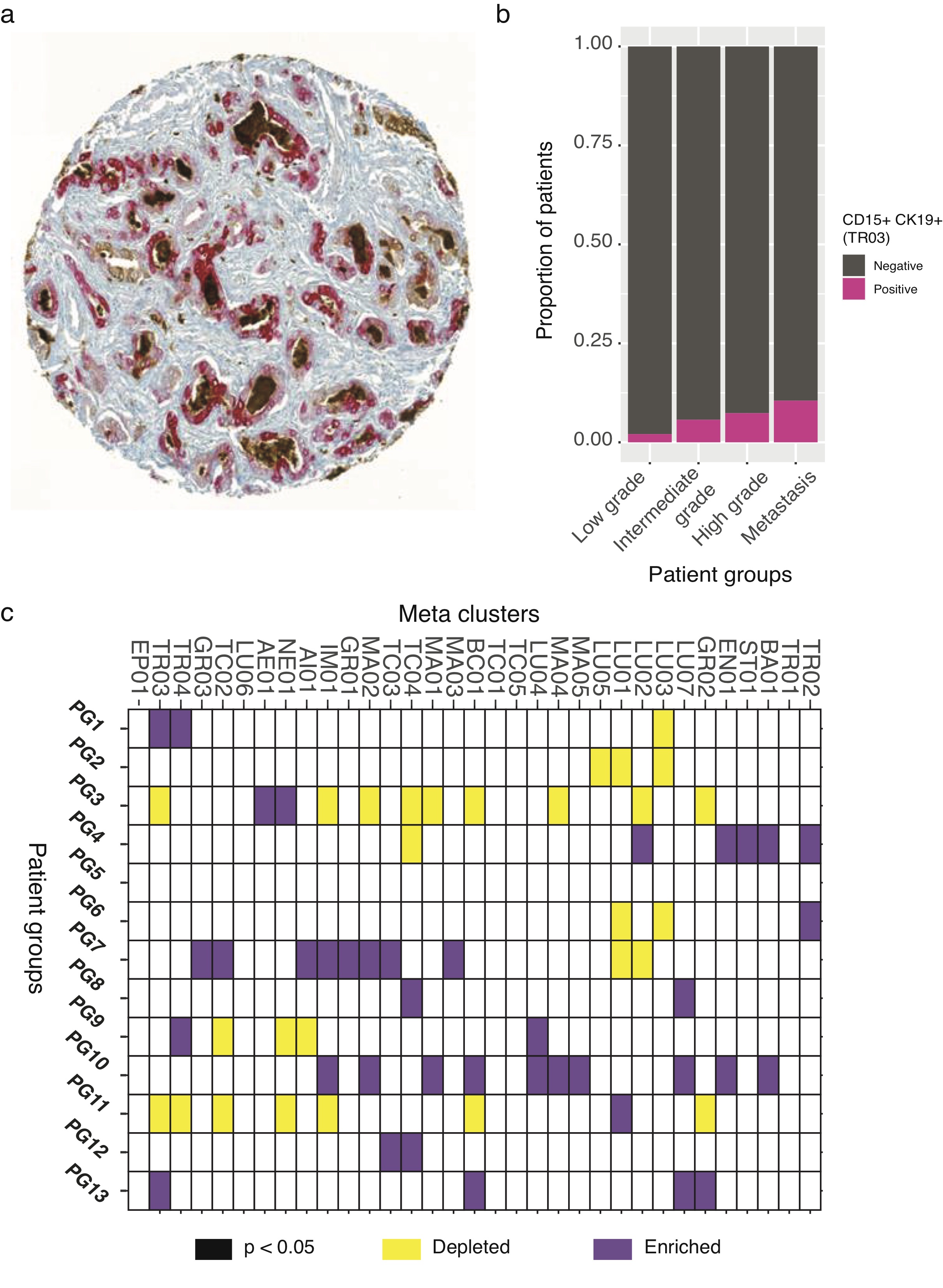
